# Variability in prior expectations explains biases in confidence reports

**DOI:** 10.1101/127399

**Authors:** Pablo Tano, Florent Meyniel, Mariano Sigman, Alejo Salles

## Abstract

Confidence in a decision is defined statistically as the probability of that decision being correct. Humans, however, display systematic confidence biases, as has been exposed in various experiments. Here, we show that these biases vanish when taking into account participants' prior expectations, which we measure independently of the confidence report. We use a wagering experiment to show that modeling subjects' choices allows for classifying individuals according to their prior biases, which fully explain from first principles the differences in their later confidence reports. Our parameter-free confidence model predicts two counterintuitive patterns for individuals with different prior beliefs: pessimists should report higher confidence than optimists, and, for the same task difficulty, the confidence of pessimists should increase with the generosity of the task. These findings show how systematic confidence biases can be simply understood as differences in prior expectations.

---

A level of confidence accompanies all of our decisions [1]. This sense of confidence can be reported explicitly, or implicitly through behavioral markers such as the amount of time willing to wait to obtain a response [2], reaction times [3], or the predisposition to wage [4] or opt-out of the task for a lower but safe reward [5]. The use of such implicit measures has shown that a sense of confidence is present even in rodents and nonhuman primates (see [6] for a review). A quantitative approach to confidence helps formalize the concept and unify its different manifestations. In statistical decision theory, the normative definition of decision confidence is the probability of the choice being correct [7–10].

Many models of how confidence emerges in the brain have been proposed, such as accumulators [2, 11–13], drift diffusion [3], and attractor dynamics [14, 15]. In these models, confidence metrics such as the difference between decision variables, post-decision evidence, or reaction time combined with evidence [3, 15, 17, 18], are interpreted as algorithmic constructions from variables available in the decision-making process [15]. Depending on the parameter values, these confidence metrics can either closely approximate the probability of the decision being correct, or show instead systematic deviations from optimality.

Recent studies, however, suggest that several aspects of confidence reports can be explained by interpreting human decision confidence as a normative readout of the probability of being correct. [16, 19, 29]. These studies are framed in the more general resurgence of rationality as a paramount human trait, which came about with the realization that probabilities are the proper language in contexts of uncertainty such as these we encounter in everyday life [20, 21, 22]. This “Bayesian Rationality” program came a long way in explaining human behavior in a wide range of higher cognitive domains, such as intuitive physics [23], intuitive psychology [24, 25], or causal inference [26]. Multimodal sensory integration remains a classic illustration of the flexibility and optimality of our inference mechanisms [27, 28].

The normative account of confidence and its supporting evidence may seem at odds with the fact that people often deviate from the statistical definition of confidence, systematically over- or underestimating their actual probability of being correct in many tasks [8, 30–39]. A given individual can be overconfident in a certain task while being underconfident in another, indicating a functional difference between the human report and the normative probability, and different individuals can variously deviate from normative confidence in a task without differing in actual performance [35,64]. Such deviations from optimality are still poorly understood. Griffin and Tversky [33], for example, exhibited cases of over- and underconfidence in intuitive judgements, and proposed that those biases arise because people do not take into account the reliability (a.k.a. weight, credence) of the evidence at hand. Since taking into account the reliability of evidence is a landmark of statistical computations, Griffin, Tversky, and several others dismissed the statistical framework as unsuitable for the understanding of confidence.

Nonetheless, there are many reasons why subjects may deviate from optimality. While it may well be the case that subjects’ confidence computation does not adhere to statistical principles, a departure from optimality might also arise even if one uses such principles but computes only approximately, or employs incorrect priors [40, 41]. Indeed, theory predicts that two Bayesian (hence, rational) agents with different prior beliefs will be relatively overconfident about the accuracy of their estimators [42], and the miscalibration between confidence reports and the probability of being correct can be explained by disagreements between the decision maker’s prior beliefs and the ones assumed by the experimenter [18]. Here, we build on this idea, adding confidence reports to a popular multi-armed bandit wagering task [49–51] to ask whether the apparent irrationality of confidence can be reconciled with the statistical account if participants’ prior expectations are taken into account.

More precisely, we encode the expectations about payoff probabilities as priors entering the Bayesian inference mechanism, and show that the between-subject variability in these priors is sufficient to explain why some people are more confident than others, why this difference is exacerbated in specific task conditions, and why they show different functional dependence on task factors. Indeed, agents can be said to be optimistic (pessimistic) if they have high (low) prior expectations, and they can have different strengths in their prior beliefs. Throughout, we use the terms optimism and pessimism to refer only to prior expectations about payoff probabilities; these expectations were uncorrelated with the general notion of optimism measured in the Life Orientation Test [65], see Discussion.

To avoid the inherent circularity of accounting for biases with priors [48], we characterized participants’ prior payoff expectations using only their gameplay strategy, studying their standpoint in the exploration-exploitation tradeoff inherent to bandit tasks. This measure of prior bias is thus defined independently of the confidence reports that we aim to explain.

Our analytic approach was the following. First, we measured subjects’ prior beliefs and established that they were an idiosyncratic trait, stable throughout the task. Then, using Bayesian probabilistic learning, we inferred subjects’ beliefs about the machines’ reward rates at the moment of the confidence report. The parameter-free model (i.e. no free parameters are needed to translate the participant’s experience into a confidence level) predicts that normative statistical confidence should be overall *lower* in optimistic subjects, and their confidence should depend differently on task difficulty. More strikingly, unlike the confidence report of an unbiased agent, pessimists’ confidence should depend less strongly on the difficulty and more strongly on the generosity of the task. Model predictions from the measured priors closely match human patterns of confidence reports.

## Results

59 participants played a modified version of the popular multi-armed bandit game, in which the subject faces several simple slot machines (two here) associated with different reward rates, and is told to maximize her reward. When an option is chosen, the result can be a success (i.e. receive a reward) or a failure (i.e. receive no reward). The machines’ reward rates were varied across different blocks, but held constant through the 16 trials comprising each block (see Figure 1). Participants repeatedly decided which machine to play, and observed whether this choice was rewarded or not at that specific trial. The machines’ reward rates were unknown to them, but they could be learnt over the course of each block.

**Figure 1.**
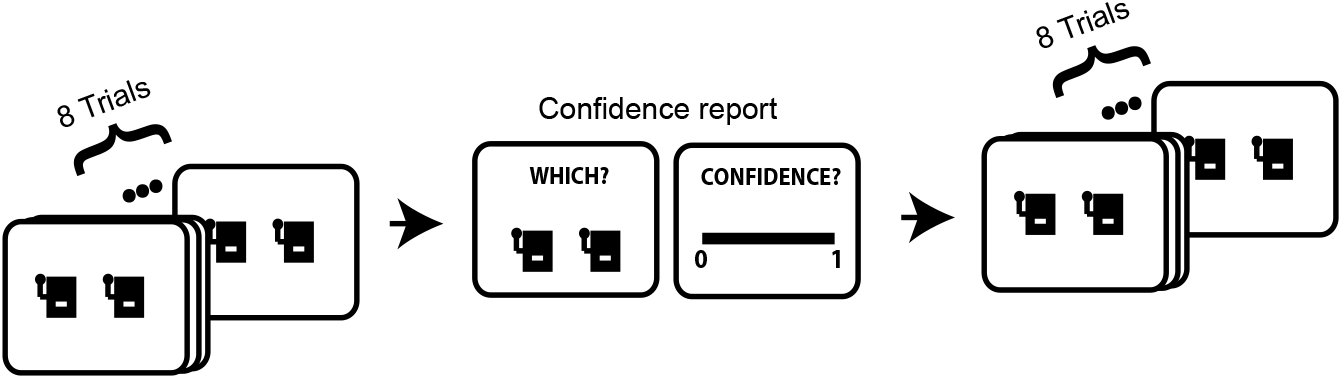
The bandit gambling task. The figure shows the structure of one experimental block. Each block consisted of 16 trials. In the middle of some of the blocks participants were required to report which machine had the higher nominal payoff and to provide a continuous report between 0 and 1 for the confidence in that decision. Each participant played 45 blocks in which confidence was reported.

From a Bayesian perspective, beliefs about the machines’ payoffs are represented with probability distributions. Observers began each block (wherein payoffs were fixed) with two identical beta prior distributions for the reward probabilities, each corresponding to one of the two machines in the experiment. We parameterize these distributions by the expected payoff rate, *b*, and the relative weight of the prior belief with respect to new observations, *w* (see *Methods*). Low values of *b* (<0.5) correspond to a pessimistic perspective, while high values (>0.5) represent a more optimistic take. The value of *w* quantifies how much the agent trusts her prior beliefs. These values (*b* and *w*) were fitted to subjects’ gameplay choices, independently of their confidence reports. Belief distributions were then subjected to bayesian updating after each machine choice made by the participant and its corresponding outcome (see *Bayesian knowledge update* in *Methods*). As shown in Fig. 4a, given a subject’s beliefs at a certain point in the block, the model’s gameplay decisions are made by taking a sample from a Normal distribution with mean *1-d* and standard deviation o. Here, the perceived difficulty *d* is equal to one minus the absolute difference between the means of the machines’ distributions at the moment of the report (machines’ payoffs differ more in easier blocks), calculated using the agent’s prior beliefs (see Eq. 2 in *Methods*).

In the middle of a block, subjects were occasionally asked to report which machine payed more, and also to indicate their confidence in that answer. We formalized this confidence report as the probability of having correctly identified the best machine, given the inferred payoff rates at the moment of the report (see Eq. 1 in *Methods* for the formal expression). Importantly, these confidence levels are read out from the learnt distributions directly, without further parameter fitting and independently of the decision-making process. In other words, the normative account of confidence in the ‘which machine is better’ decision depends only on the answer to this question and on the posterior distributions at the moment of the report, but not on the decisions that gave rise to those distributions.

### Consistency of the prior bias

We first model gameplay behavior ignoring the confidence report. For each subject, we fitted the model to actual choices using Maximum Likelihood Estimation with varying *b, w* and *σ* (MLE, see Maximum likelihood fitting in Methods). Bayesian Model Comparison indicated there was a probability *xp>0.99* (exceedance probability, see Model comparison in *Methods*) that the model in which only *b* was varied was better than a model that varied any combination of *b*, *w* and *σ*. This indicates that the additional complexity that is introduced in the model by adding other free parameters besides *b* does not increase the likelihood of the data significantly. Furthermore, varying *b* alone is sufficient for the model to show a range of behaviors that covers the entire region displayed by human participants (see Fig. 2). This does not happen when varying *w* or a alone. Therefore we settle on this simple model, varying only *b* across subjects, and fixing *w* and a to their ML global estimate at the group level (20 and 0.05 respectively).

**Figure 2.**
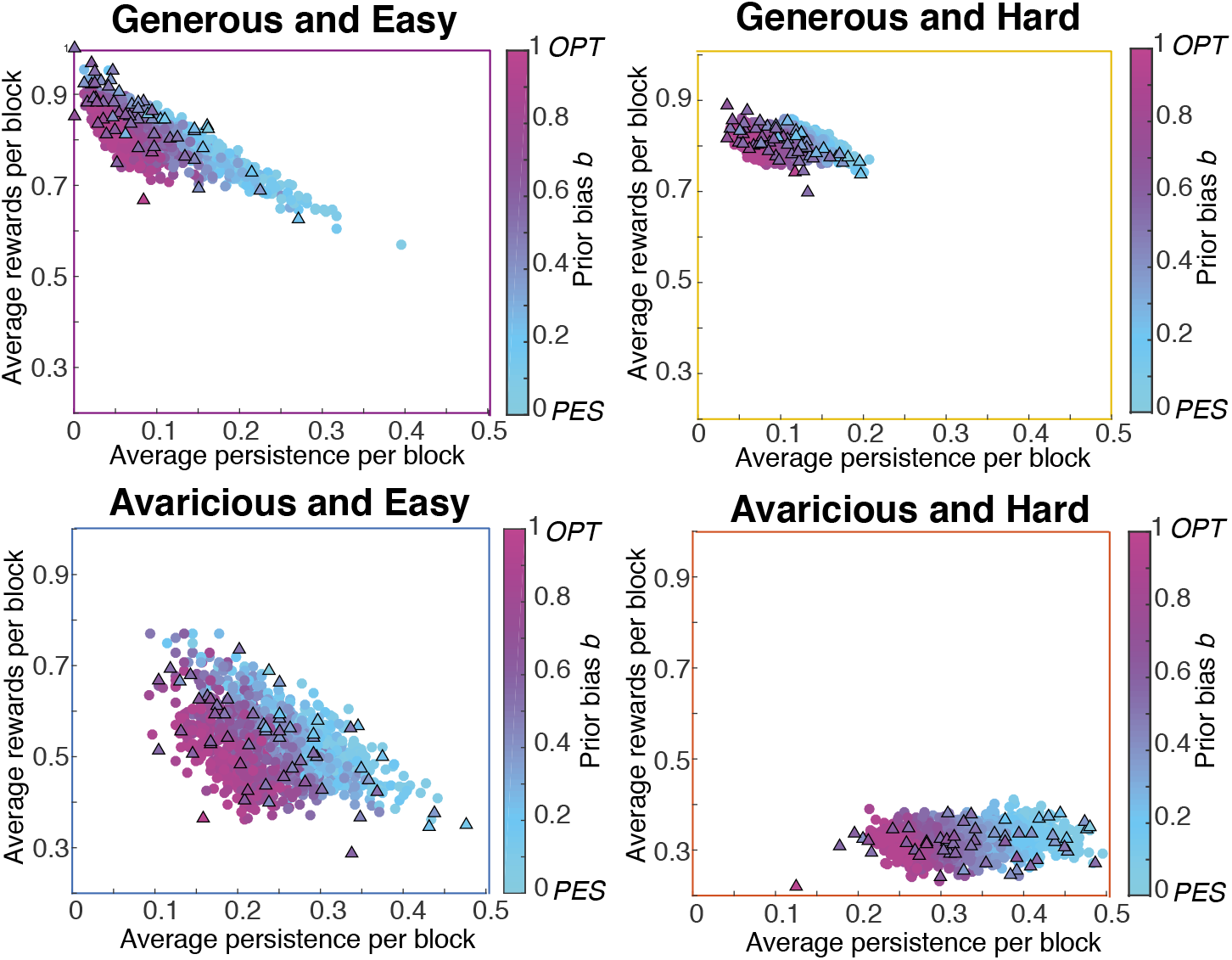
The human range of choice behaviors is accounted for solely by the prior optimism level. “Average persistence per block” corresponds to the proportion of trials in which a machine is chosen immediately after a no reward trial in that machine. “Average reward per block” is the proportion of trials in which a reward was obtained. Each of these panels shows behavior averaged over a set of blocks, divided according to their generosity and difficulty. Participants are shown with black-edged triangles. The color within each triangle corresponds to the MLE *b* value fitted for each participant using all blocks (see Methods). Each participant has the same color in all panels. The coloured points correspond to 600 runs of the model. In each run, *w* and a were fixed to their group-level MLE fitted values (20 and 0.05 respectively) and *b* varied between 0.05 and 0.95. Since *w* and *σ* were kept fixed to their group-level, best-fitting values, the entire variation of human behavior is accounted for solely by the variation of the prior mean *b*. Conditions are matched to the curves in Fig. 3a by the coloured frames.

The prior bias *b*, which captures the level of optimism, was fit individually for each participant using a MLE approach (see Maximum Likelihood fitting in *Methods*). To test the idiosyncratic nature of this prior, we verified that it consistently impacts behavior across different conditions. More precisely, we split all blocks in Generous and Easy, Generous and Hard, Avaricious and Easy, and Avaricious and Hard by median splitting the blocks according to their unbiased difficulty (one minus the difference between machines’ mean payoff, calculated with an unbiased prior, see Eq. 2) and unbiased generosity (average between machines’ mean payoff, calculated with an unbiased prior). For each subject, we fitted the ML value of *b* independently in the four conditions (Fig. 3a). We then performed a cross-validation test by comparing the fitted value from all data in three of the four conditions with the fitted value from the left-out condition (Fig. 3b). By choosing any of the four conditions as the left-out condition, we have four points for each subject. For all points from all subjects, the correlation was *ρ*=0.65 (p<0.0001), and the mean squared error of predicting each subject’s *b* value from the left out category using the average *b* value of the other three categories was 0.02, amounting to an absolute error of 14%. Finally, if we fit the *b* value of each participant independently in each block (as shown in Figure S4, see Supplementary Information) we see that for most participants the fitted value remains in either pessimistic or optimistic levels during the entire experiment. This establishes the value of *b* as an idiosyncratic, stable individual trait.

**Figure 3.**
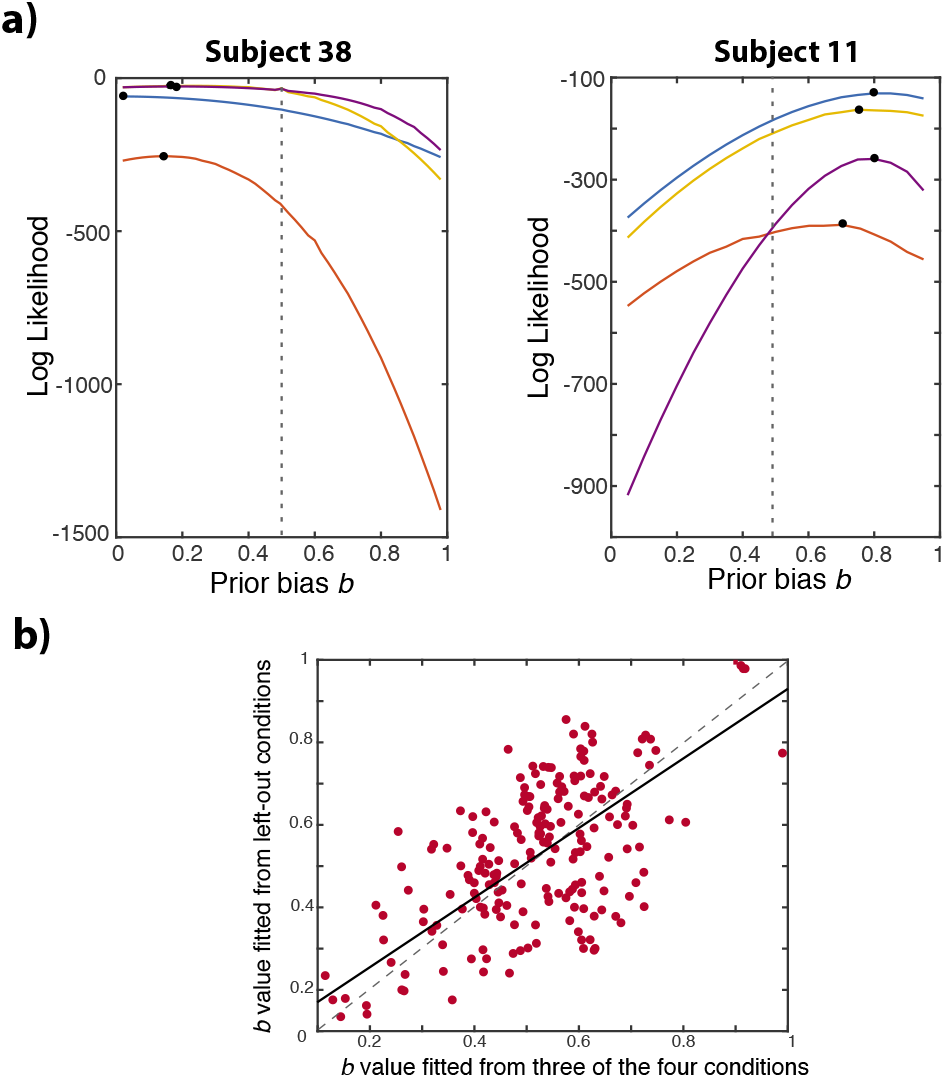
Prior biases adjusted from gameplay are idiosyncratic traits, stable across conditions. **a)** Blocks were divided in four conditions according to their generosity and difficulty. Here, we show two example participants, a pessimistic one (left) and an optimistic one (right). The likelihood of choices under each value of the prior bias *b* was calculated independently in each of the four conditions, represented by the four lines in each panel, with colours matching the frames in Fig. 2. Black dots show the MLE of *b* in each condition. From these two panels we can see that consistent values of *b* are fitted for the same participant across the four different conditions: for each subject, the black dots are in similar *b* values. The panels for all participants are shown in Figure S3 (see Supplementary Information). **b)** The consistency of fitted *b* values for all participants can be quantified in a cross-validation test. Each point corresponds to the average MLE of the *b* value fitted for one subject in three of the four conditions (horizontal axis) compared with the value fitted independently for the left-out condition (vertical axis). The correlation coefficient between the two values is ρ=0.65 (p<0.0001). The black line is a linear fit (slope is 0.8 (t=13), R^2^=0.42), close to the identity line (dashed). These results indicate that the *b* value is an idiosyncratic trait that can consistently describe participants’ behavior in any condition.

### Pessimists report higher confidence than optimists

An important finding is that optimists and pessimists answered the question “Which machine pays more?” prompted in the middle of each block with similar accuracy, computed as the fraction of the blocks in which the machine with the higher real payoff was correctly identified (0.70±0.09 s.d. for pessimists and 0.70±0.06 s.d. for optimists, a two-sample t-test gives t(52)=0.2, p=0.8). Therefore, differences in the reported confidence levels cannot be accounted for by a difference in actual performance.

In Figures 4, 5 and 6 we compare participants’ confidence reports to the probability of having made the correct decision, calculated with an unbiased prior (indicated as ‘No-bias model’) and calculated with participants’ priors as adjusted from gameplay (indicated as ‘Model’). That is, for each participant and block, we take the history of successes and failures *experienced by the participant* and compute the posterior beliefs that the participant should have at the moment of the report, using an unbiased prior as the initial distribution (no-bias model) or using their adjusted priors from gameplay as the initial distribution (model). Then, given these posterior beliefs, we compute the probability that the decision *made by the participant* to the ‘which machine is better’ question was correct. Finally, we compare this probability under the two models with the participant’s report (indicated as ‘Humans’).

The model predicts that pessimists (low *b*) should have higher confidence than optimists (high *b*) particularly when one option is exploited and the other one is left unexplored, which is often the case in easy blocks or in blocks with high generosity. Indeed, if we separate participants according to their *b* value being greater (N=25) or smaller (N=29) than 0.5, pessimists and optimists confidence levels in high generosity conditions (generosity>0.5, median split) were respectively 0.78±0.02 s.e.m. and 0.70±0.03 s.e.m. (two-sample t-test: t=2, p<0.05; N=3 participants had *b*=0.5 and hence left out of this categorization). Although it may appear counter-intuitive that pessimistic people are more confident, this can be easily understood: expecting less, whenever they find a machine that pays somewhat well, they are very confident that this machine is indeed better than the other one. This in turn is amplified because better machines are played more, so beliefs about the unexplored machine remain dominated by the pessimistic prior. Optimists on the other hand expect more, so playing a good machine does not separate its reward distribution that much from the prior, and thus confidence is lower. Following this intuition, one could expect optimists to report higher confidence than pessimists when a bad option is chosen repeatedly, so the effect should fade away on average. In practice, however, this situation does not happen: only good options are chosen repeatedly, leading to pessimists reporting higher confidence than optimists. This effect is illustrated in Fig. 4a.

In our framework, there is a continuum between optimism and pessimism, quantified by the values of prior bias *b*. The model predicts that subjects who are all the more optimistic (larger *b*) should overall have lower confidence levels. This is shown in Fig. 4b (middle): the average confidence level should decrease with the value of the prior bias *b*, as quantified by the negative slope of the linear fit (-0.36±0.02 s.e.m.). Indeed, we show in Figure 4b (left) that the average confidence reported by participants depends on their adjusted prior from gameplay in the expected way. The correlation between human average reported confidence and the prior bias *b* value is *ρ*=-0.37 (p=0.004), and the slope of the linear fit is -0.27±0.01 s.e.m. (t(55)=3, p<0.005).

**Figure 4.**
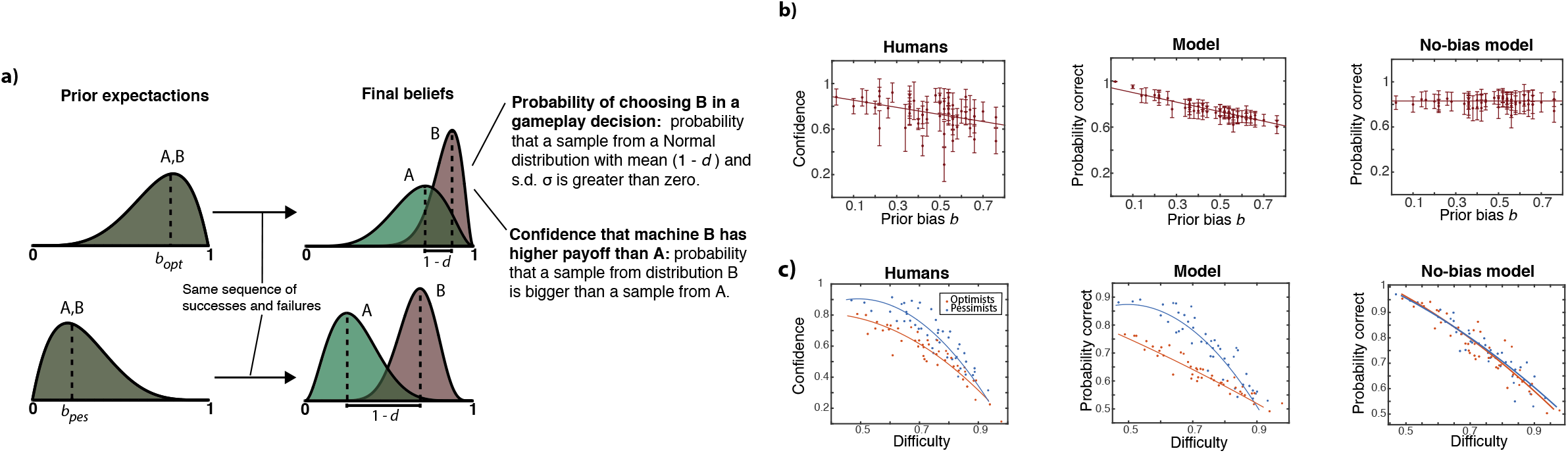
Pessimists report higher confidence than optimists. **a)** After experiencing the same sequence of successes and failures (6 successes and 0 failures for A and 0 successes and 2 failures for B in this example), confidence from a subject with low prior expectations (bottom, low *b*) should be higher than confidence from a subject with high prior expectations (top, high *b*). This happens because, expecting less, whenever pessimists choose repeatedly an option that is generously rewarded (B), the distribution from the option left unexplored (A) remains in lower values than for optimists, yielding a higher probability of being correct for pessimists than optimists (lower perceived difficulty *d*). **b)** ‘Humans’ shows humans’ confidence confidence report. ‘Model’ corresponds to the probability correct calculated using participants’ experience at the moment of the report as perceived by a rational agent with a prior equal to participants’ priors adjusted from gameplay. ‘No-bias model’ corresponds to the probability correct calculated using participants’ experience at the moment of the report as perceived by a rational agent with an unbiased, non-informative prior. Each point represents the average confidence report of a subject in all blocks (Left), or the average probability of being correct in all blocks (Middle and Right). Subjects are classified according to their prior bias *b* adjusted from gameplay (*x*-axis). In these plots we analyze only high generosity blocks (*Generosity* > 0.5), where the expected difference between agents with different *b* values is bigger. Error bars are s.d. across all blocks. **c)** We divide participants into pessimists (blue, N=25) and optimists (orange, N=29) according to their adjusted *b* value from gameplay being strictly smaller or greater than ½. Each point represents the average confidence (or probability of being correct) level of all subjects in a given block, classified according to the block difficulty (all blocks are shown). Error bars are omitted for clarity (average s.d. is 0.15).

Importantly, we show in Figure 4b (right) that the average probability of having made the correct decision computed with an unbiased prior is similar for all subjects (linear fit slope equal to 0.00±0.02 s.e.m.). This means that, on average, all participants experienced equivalent histories of successes and failures until the moment of the report: from an unbiased perspective, they should all share similar beliefs and have no reason to report different confidence levels. Finally, we note that the average accuracy to the ‘which’ question in high generosity situations (>0.5) does not correlate significantly with the value of *b* (t(55)=0.93, p=0.35). Therefore, the variation of the confidence level for different *b* values observed in Fig. 4b is not due to differences in actual performance.

### Different effects of difficulty on confidence in pessimists and optimists

As suggested in Figure 4a, in order for the asymmetry between the confidence level of optimists and pessimists to appear, one of the options should be left rather unexplored, and therefore more influenced by its prior position. In situations in which both options are chosen frequently, prior beliefs fade and the initial position of the distributions is irrelevant. Going back to the grouping of participants into pessimists and optimists according to their adjusted *b* value, we show in Fig. 4c (middle) that in theory, the expected value of the difference between optimists and pessimists is bigger in easier blocks, where the option is often left unexplored and therefore more influenced by its prior position (this situation is also often in high generosity blocks). In hard blocks, contrastingly, both options are often explored, and prior beliefs fade, so optimistic and pessimistic agents should report similar confidence. This is precisely the pattern we find in humans (Fig. 4c, Left). We quantified this effect by performing a regression analysis for all human responses in all blocks. The linear model was of the form *c_s,k_ = β_0_ + β_1_d_k_ + β_2_b_s_d_k_* where *c_s,k_* is the reported confidence by participant *s*, whose prior bias is *b_s_*, in a block with difficulty *d_k_*. The adjusted value of *β*_2_ = -0.24 ± 0.04 (t=6.62, p<10^-10^) reveals a significant interaction between task difficulty and prior bias on the reported confidence level. The model that accounts for participants’ priors shows *β*_2_ = -0.33 ± 0.01 ( t=25, p=0).

Importantly, the participants’ probability of being correct that is calculated with an unbiased prior is mostly the same for optimists and pessimists (Fig. 4c Right), indicating that gameplay experience was similar for both groups: without considering agents’ prior beliefs, both groups should have reported similar confidence values throughout, and there is no reason to believe their average confidence level should depend on block difficulty in different ways. In the regression analysis, the no-bias model showed a small interaction between *b* and *d* in the opposite direction to the one seen in humans: *β*_2_ = 0.06 ± 0.01 (t=4, p<10^-5^). Finally, we study accuracy to the ‘which’ question in a regression analysis of the form *a_s,k_ = β_0_ + β_2_d_k_ + β_2_b_k_d_k_*, where *a_s,k_* is equal to 1 if the machine with the higher real payoff was correctly identified by participant *s* in block *k*, and 0 if not. Besides an obvious dependence on difficulty (*β_1_ = -0.66 ± 0.06 (t=10, p=0)*), we find no significant dependence of accuracy on the interaction between the prior bias *b* and block difficulty: *β*_2_ = -0.07 ± 0.07 (t=1, p=0.34), indicating that the different dependence of confidence on task difficulty for participants with different *b* is *not* due to different accuracy in the ‘which’ question among these participants.

### Pessimistic confidence should increase with the generosity of the task

To compute task difficulty, we must assume *some* prior beliefs (see Eq. 4 in Methods). From an experimenter point of view, the more ‘reasonable’ choice is the non-informative, unbiased prior (*b*=0.5 and *w*=2). When difficulty and generosity are computed in this way, the confidence report of an unbiased agent depends mostly on the difficulty of the task. However, for a pessimistic agent, the reported confidence depends not only on the difficulty of the task but also on its generosity: for the same difficulty, normative pessimistic confidence increases with the generosity of the task.

To explain the interaction between task factors and subjects’ prior beliefs, we compare how these beliefs update for a pessimistic agent after two different situations with the same difficulty (0.38) but different generosity (0.44 and 0.56) in Fig. 5a (left column). The key to understanding this effect is that if the observations of a pessimistic agent align with her pessimistic prior, then the prior distribution will not change its position, but will only become more peaked. The red distribution (in Fig. 5a) for the pessimistic agent is therefore practically at the same location after a few failures (situation 2) than after several more failures (situation 1, compare bottom and top rows). In contrast, observing many successes is unexpected for a pessimistic prior. This is illustrated in the green option, which returns a few successes (and no failures) in situation 1, and a lot of successes (and no failures) in situation 2. Instead of increasing in height, the green distribution moves towards more optimistic values as more successes are encountered.

**Figure 5.**
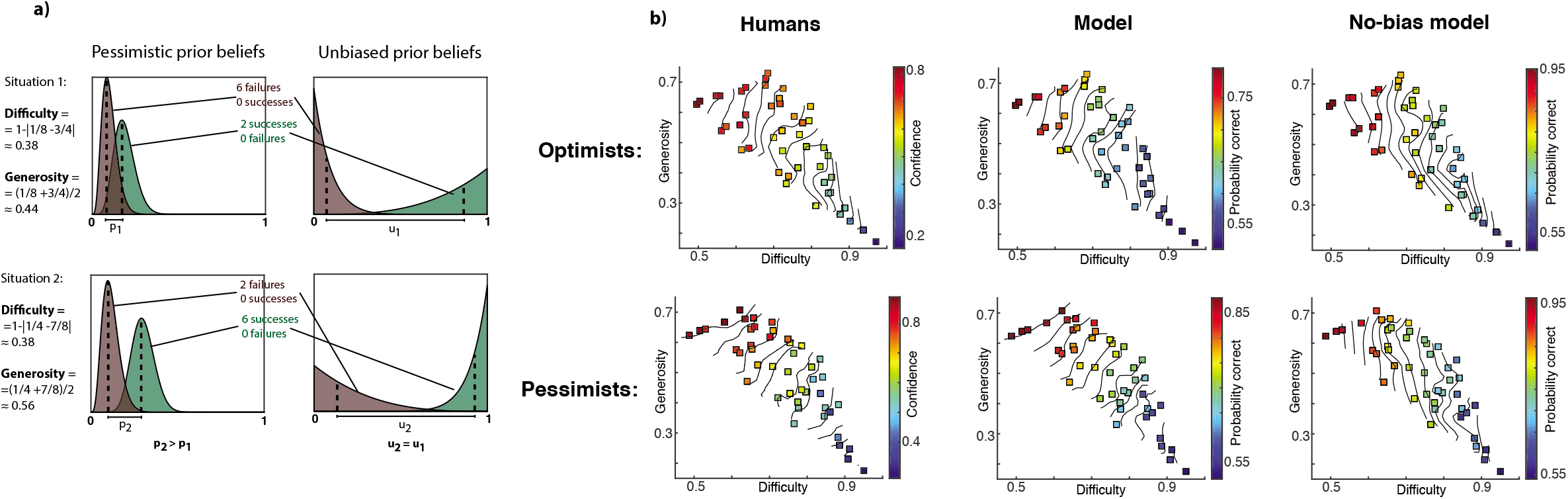
For the same task difficulty, pessimistic confidence increases with generosity. **a)** Each row shows the final beliefs that a pessimistic (left) or an unbiased (right) agent should have after receiving the experience corresponding to that situation. In situation 1 the red option returned 6 failures and 0 successes, and the green option returned 2 successes and 0 failures. The difficulty and generosity of each condition is indicated on the left margin. **b)** Humans are classified as optimistic or pessimistic according to their prior bias *b* adjusted from gameplay being bigger or smaller than respectively. As in previous figure, ‘Model’ computes probability correct using subjects’ experience as perceived with their prior adjusted from gameplay. ‘No-bias model’ that computes probability correct using subjects’ experience as perceived with an unbiased, uniform prior (*b*=0.5, *w*=2). Each square shows the average confidence reported by all optimistic (top row) and pessimistic (bottom row) subjects on the experimental block with that generosity (y-axis) and difficulty (x-axis). Both difficulty and generosity were calculated with an unbiased prior. Black lines are isoconfidence lines.

In sum, data that is incompatible with an agent’s strong prior beliefs produce bigger changes in those beliefs than data compatible with them. The two situations from Figure 5a, left column, which have the same final difficulty but differ in generosity, will show virtually the same distribution for the red option, but the green distribution will end up around more optimistic beliefs for the high generosity situation than for the low generosity one. In other words, situations 1 and 2 are not symmetric: the green distribution moves more to the right in situation 2 than the red distribution moves to the left in situation 1, and the separation of the two distributions will be bigger in the high generosity situation (p2>p1), thus leading to higher confidence.

The unbiased agent, on the other hand, will display symmetrical beliefs in the two situations (Fig. 5A, right column), with the red distribution moving as much to the left in situation 1 as the green distribution moves to the right in situation 2 (u2=u1). The distributions are equally separated after either of the two situations, leading to the same confidence. The fact that pessimistic confidence should significantly increase with generosity for the same task difficulty means that the isoconfidence lines of pessimistic agents in Figure 5b (middle) should not be vertical like for the unbiased agent (right), but instead show a positive slope, indicating that confidence decreases with difficulty but also increases with generosity.

Following this intuition, we would expect optimistic agents to report, for the same difficulty, higher confidence in low generosity situations than in high generosity situations. For this to happen, optimistic players should repeatedly choose an option that does not give rewards. As mentioned before, this does not occur in practice. Instead, in low generosity situations both options are chosen evenly and the reported confidence of an optimistic agent is more in line with the real, unbiased difficulty of the task (i.e. more vertical isoconfidence lines in low generosity conditions).

Independently of the intuition we can build for this phenomena, human results, shown in Fig. 5b, are striking: optimists and pessimists show very different confidence patterns, as predicted by the model. Importantly, the no-bias model is unable to capture the strong biases we see on pessimists’ isoconfidence lines. Without accounting for participants’ prior beliefs, and instead computing confidence using unbiased prior beliefs, there would be no reason to believe that, for a fixed difficulty, pessimistic confidence should increase with the generosity of the task. Therefore, pessimistic participants would be judged as having a very strong confidence bias with respect to the rational model if an unbiased prior is assumed for all participants. However, this bias is precisely what it is expected when we account for their strong pessimistic prior adjusted from gameplay. We quantified this by performing a regression analysis of the form of the form *c_s,k_ = β_0_ + β_1_d_k_ + β_2_g_k_d_k_ + β_3_b_s_d_k_ + β_4_b_s_g_k_d_k_* where *c_s,k_* is the confidence reported by participant *s* whose prior bias is *b_s_*, in a block with difficulty *d_k_* and generosity *g_k_*. The regression shows a significant effect of difficulty (*β*_1_= 0.9±0.2, t=4, p<10^-5^) and an interaction between difficulty and generosity (*β*_2_= - 1.3±0.2 t=6.8, p<10^-11^). As expected from the results of the previous section, there was a significant interaction between the effect of difficulty and the prior bias of participants’ (*β*_3_=-0.6±0.2, t=3.2, p=0.001). More importantly, we also find a significant triple interaction between subjects’ prior bias, generosity and difficulty of the task (*β*_4_=0.5±0.2, t=2, p=0.04), indicating that the prior bias *b* affects the way in which difficulty interacts with generosity. The model that accounts for participants’ priors shows significant interaction between *d* and *b*: *β*_2_=-0.62±0.07 (t=8.9,p=0); and a significant triple interaction between *b, d* and *g: β*_4_=0.35±0.08 (t=4,p<10^-4^). The no-bias model, on the other hand, predicts no significant interaction between *b* and *d: β*_2_=0.11±0.07 (t=1.5,p=0.13); and no significant triple interaction between *b, d* and *g: β*_4_=-0.05±0.08 (t=0.5,p=0.55), as suggested by the vertical isoconfidence lines. Similarly, the accuracy to the ‘which’ question does not depend significantly on the interaction between *b, d* (t=1, p=0.2) or between *b, d* and *g* (t=1.4, p=0.15), indicating that the different inclination of isoconfidence lines for participants with varying *b* is not due to a different dependence of accuracy on task difficulty and generosity.

The differences between the no-bias model and pessimists’ confidence reports shown in Fig. 5b are strong enough to cause the confidence reports of all subjects, pessimists and optimists lumped together, to strongly differ from the ones of the model with no bias. This is shown in Fig. 6. Analyzing the report of all subjects, we see that humans show isoconfidence lines markedly different to the ones of the no-bias model, but their biases are precisely those expected by the model that accounts for their prior beliefs fitted from gameplay. Again, without accounting for participants’ prior beliefs, there is no reason to think that human confidence should increase with the generosity of the task, and humans would be judged having a systematic confidence bias because of their failure to display vertical isoconfidence lines. However, this bias is precisely what is expected when we account for participants’ priors adjusted separately from gameplay.

**Figure 6.**
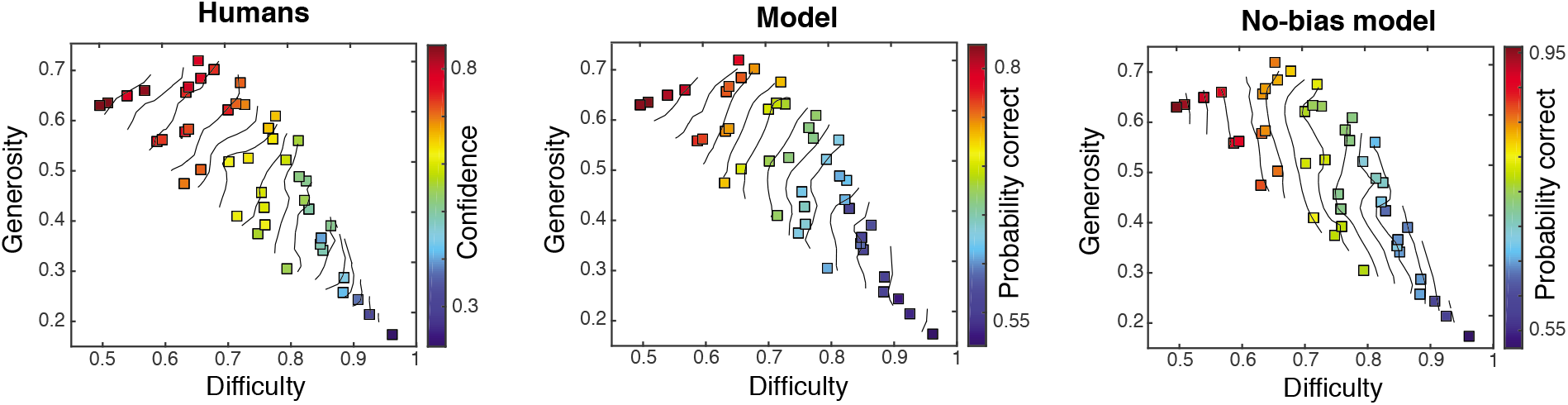
Human biases on isoconfidence lines are explained by the prior optimism level measured in gameplay. Legend as in previous figure. Squares show the average confidence report of all subjects, without separating pessimists from optimists.

### Calibration of confidence

As is elegantly explained in [29], isoconfidence lines are a robust characterization of the human sense of confidence. Unlike the study of confidence calibration (i.e. comparing the numerical probability of being correct with the numerical value of the confidence report), isoconfidence lines do not change if we rescale the confidence reports with any monotonic function and they are independent of the particular way in which participants may be using the confidence bar to report it. However, we can still analyze the effect that accounting for participants’ priors has on confidence calibration. As shown in Fig. S1 (see Supplementary Information) the difference between the numerical confidence report given by humans (in a scale from 0 to 1 indicated by the region of the confidence bar) and the probability of being correct calculated using the model that account for participants’ priors is similar in all blocks, humans being underconfident by a similar amount in every block. If we instead calculate probability correct using human experience at the moment of the report starting from an unbiased prior, humans report a higher confidence value with respect to the probability correct in generous blocks than in avaricious blocks, this is, the magnitude of the underconfidence bias is not the same in all conditions, but depends on their generosity.

## Discussion

We presented a two-armed bandit experiment in which participants maximize their rewards by playing machines with unknown payoff rates. Participants explored the machines and were occasionally asked to report their confidence about knowing which machine payed more. We first showed that each participant can be characterized with a consistent prior bias according to their choice behavior during the game, independently of their confidence reports. The statistical modeling of confidence predicts that subjects classified as pessimists should report higher confidence than subjects classified as optimists, even though all subjects had similar accuracy and similar evidence available to correctly identify the best machine. The model also predicts that this difference between optimists’ and pessimists’ confidence should be exacerbated in high generosity and easy situations, and, for the same difficulty level, pessimistic confidence should increase with the generosity of the task. Participants’ behavior conformed to all these predictions. Our results indicate the variability of confidence reports across participants are expected when taking into account their different prior beliefs.

Importantly, our predictions are robust to the specific choice of the form of the confidence readout from the belief distributions at the moment of the report. While we reported results using the normative statistical approach (i.e. Eq. 1 in *Methods*), similar effects can be predicted when computing confidence with other metrics like the distance between the means, or the overlapping area between both distributions, to name a few. The critical aspect of our model is not the particular form of the confidence readout from the belief distributions, but that the estimation of those underlying distributions should take into account the participants’ prior expectations. Additionally, since prior expectations are estimated from independent data (the gameplay), the model for confidence has no adjustable parameters: its predictions are inescapable.

Taking into account participants’ prior beliefs is key to evaluating the rationality of their behavior in our task. Such an approach should nevertheless avoid two pitfalls. The first is circularity: *assuming* a difference by appealing to priors rather than *explaining* it, which can be a limitation of Bayesian models. Indeed, in principle, any behavior can be accounted for by a particular set of prior beliefs together with a decision rule [48, 54]. The second pitfall is overfitting: improving the goodness-of-fit by resorting to more free parameters. Our approach avoids these pitfalls by using a parsimonious, general Bayesian model that accounts for both choice and confidence. Indeed, we assumed that these two aspects of behavior are exclusively affected by a single set of prior beliefs and the pattern of rewards experienced in the game. Similarly to cross-validation methods, where data are divided into training and test sets [55] in order to limit the complexity of the model (i.e. the number of free parameters), our experimental design is divided into “training” and “test” tasks (machine choices and confidence report, respectively). Behavior in the training task is used to learn -and pin up- the prior parameters for each individual. If the resulting parameter-free model for the test task predicts human data accurately, it is likely to generalize well to future data, limiting the risk of overfitting [56].

A direct comparison of confidence reports between subjects can be achieved by a direct matching between the numerical probability of being correct and the human confidence report, i.e. the calibration of confidence [8]. However, this comparison requires that subjects report confidence on the same absolute scale, which is untestable in practice, forcing the experimenter to choose a particular reparametrization function for mapping the experimental confidence measurement into a value between 0 and 1. Actually, every confidence calibration result, here and elsewhere in the literature, depends on the chosen reparameterization function, and on the way in which different participants use the confidence bar. For example, if in our experiment we would interpret the confidence bar as going between 0.5 and 1 instead of between 0 and 1, humans would be judged as being *over*confident on average instead of underconfident. Indeed, Aitchison *et. al*. [29] argued that when confidence is displayed along a one-dimensional quantity (e.g. in [19]) there is always the freedom of reparameterizing confidence reports by a monotonic function, which in one dimension would allow us to trivially explain any observed pattern by a suitable such transformation. When plotting the results along two independent variables (e.g. confidence vs. generosity and difficulty) this reparameterization is no longer possible, and the arising pattern of isoconfidence lines in the plane is now a robust indicator of the participants’ behavior, independent of the way in which participants use the confidence bar. Specifically, the two dimensional representation only analyzes the transformation of incoming data into an internal representation from which confidence is readout as a continuous variable, separating it from the mapping of this continuous variable onto some external scale in order to report it, in which humans do not seem to be optimal [8, 35]. The interactions we found between task difficulty, generosity, and the prior bias (Figs. 5 and 6) are therefore robust and immune to the above mentioned issues regarding the calibration, readout and re-scaling of confidence reports. The only requirement for this analysis is that each participant uses the confidence bar in a way that is consistent across trials. Note that if we add the restriction of choosing the same (monotonically increasing) rescaling function for all participants, pessimists would report higher confidence than optimists and this asymmetry would be bigger in easier blocks independently of which function is chosen.

Our results indicate that participants’ prior expectations about reward rates in the task constitute an idiosyncratic trait, stable across task conditions. This naturally begs the question whether it can be related to a more general notion of optimism: are optimists in the task more likely to be optimistic in real-life situations? Although appealing, we don’t have the evidence to support such hypothesis. The Life Orientation Test (LOT) [65] was available for only a small subset of the subjects (n=11), and we found no correlation between our prior bias assignment and optimism as measured by the LOT score (n=11, r=0.2, p=0.54). We acknowledge that there may quite possibly be several different types of optimism, some pertaining to prior expectations (as here), others to the way we learn from positive and negative outcomes [43–47] or how we foresee the future [66], and yet others characterizing situations rather than individuals [67].

Variability in prior expectations may also explain confidence biases in other contexts. For example, it was shown in [18] that the hard-easy effect [58] (over-confidence for hard questions and underconfidence for easy ones) is expected when participants have a strong prior for the difficulty of each block arising from their experience in prior blocks. In perceptual tasks, in which the agent accumulates evidence about an input signal in order to make a decision, agents differing in their prior beliefs display different confidence biases [42]. For clarity, we compare the beliefs of a biased agent with respect to the beliefs of an unbiased agent. In this context, the hard-easy effect is expected for agents whose prior beliefs are closer to the actual signal in easy trials than in hard trials. More strikingly, subjects with prior expectations about the signal variance which differ from the actual variance will display an endogenously generated confidence bias: as Bayesian-rational agents put more weight on more precise information, the evidence with overestimated precision becomes excessively weighted in further processing [42, 57]. Indeed, variability on prior uncertainties can explain systematic confidence biases in various contexts; we can revisit the classic result of Tversky and Griffin [33] in much the same way. In their experiment, subjects’ isoconfidence lines were less dependent of the number of coin tosses (i.e. weight) and more dependent on the average result of the tosses (i.e. strength) than those of a model with no a priori uncertainty (see Supplementary Information for details). Simply allowing for uncertainty in the prior lets us reproduce human confidence patterns (see Fig. S2 in Supplementary Information). Of course, theory alone is not enough to judge participants’ systematic confidence biases, so we cannot claim that in their experiment participants were using a rational strategy but with incorrect priors. As was done in our study, it is also necessary to actually measure participants’ prior beliefs and then analyze whether the observed behavior matches that of a rational agent with those beliefs.

In this paper, we have experimentally shown how the biases apparent when comparing human confidence reports with the probability of being correct calculated using an unbiased prior disappear when we use participants’ priors instead. Importantly, we are not claiming that participants’ prior beliefs are optimal, instead, our claim is that, given the prior, humans update their beliefs in an approximately optimal manner, and confidence is then readout from the appropriate posterior distribution. We showed that prior beliefs can be major determinants in the analysis of rationality in the human sense of confidence, and seemingly reasonable (but erroneous) assumptions like a uniform prior would be in our experiment, can lead to serious mistakes when judging rationality. Our results suggest that, whenever possible, future studies of rationality should account for participants real priors instead of assuming them. This may be of interest to researchers, or even practitioners, who use confidence reports as diagnostic features of our metacognitive abilities, i.e. the ability to evaluate our own state of belief. Indeed, variations in confidence levels across individuals exposed to the very same situation and having the same actual performance typically suggest that the evaluation of confidence is dysfunctional in some subjects (e.g. confidence increasing with task generosity for the same task difficulty for some subjects). While this may indeed be the case [60] our results raise a cautionary note. One needs to first monitor the state of belief, which may differ between subjects, to then assess variations in confidence levels.

Although confidence reports have traveled a winding road in the psychology and neuroscience literature, recent work is settling in on the statistically normative account of confidence [9, 19]. This study contributes to this view, showing how systematic confidence biases can be simply understood as differences in prior expectations. This further fuels the view of humans as rational animals, and signs off another success of the Bayesian rationality program [59].

## Methods

### Experiments

Participants were informed that they were going to play a series of unrelated blocks in each of which the payoff of the machines was unknown but fixed, and their aim was to maximize the total reward in order to win a monetary prize. Each block comprised 16 trials, blocks were clearly delineated from one another by pauses, and there was a message reminding participants that separate blocks were independent from one another. The task instructions were in written format, a translation can be found in the link provided in the *Data Availability* section (original language is Spanish). Participants read the instructions and then completed 6 demonstration trials before the task, accompanied by the experimenter, to get acquainted with the graphical interface.

The data collection was divided in two studies. The experiment in Study #1 comprised three type of blocks. In the ‘confidence’ blocks (see Fig 1) participants were asked, in the middle of the gameplay to report which machine pays more and their confidence in their answer to that question. Some blocks were played uninterrupted (‘no report’ blocks) and in some other blocks participants were asked which machine is better but without a confidence report (‘which’ blocks). The experiment in Study #2 comprised two types of blocks: the standard ‘confidence’ blocks, and some ‘distractor’ block similar to the ‘confidence’ blocks excepted that the confidence question was replaced by a simple arithmetic operation that took a time approximately equal to the time of the confidence report. Experiments were designed in this way in order to also test for the effect of reporting confidence. In this paper, we focus on confidence levels themselves and therefore on the ‘confidence’ blocks.

The nominal pairs of reward rates were exhaustively drawn from a predefined list in random order and assigned to each block, where each rate in the pair was again randomly assigned to a machine. This process guaranteed that all participants played the same set of blocks and that there was no correlation between the payoffs of the two machines along the game for any participant (correlation coefficients were smaller than 0.23 and p>0.05 for all subjects). The list of pairs of rewards rates are all possible 45 pairs formed from the set {0, 0.125, 0.25, 0.375, 0.5, 0.625, 0.75, 0.875, 1}.

The task was designed and implemented in Python using the PyGame library [61], and lasted around 45 minutes (135 blocks set up, study #1) and 25 minutes (70 blocks set up, study #2) during which the participant was left alone in a quiet room.

Average performance did not show a significant decay during the task (correlation between block number and average human reward is ρ=0.01 (p=0.82)), so that all blocks were analyzed irrespective of the block number.

### Participants

A total of 59 nonpaid participants from the general population between 20 and 55 years old were recruited (28±7 s.d. years old, 24 females) from our’s laboratory subject pool. Two participants were excluded from the analysis for obtaining a total reward consistent with random play. In study #1, 17 participants completed 45 ‘confidence’ blocks, 45 ‘which’ block and 45 ‘no-report’ blocks. In study #2, 42 participants completed 45 ‘confidence’ blocks, 15 ‘distractor’ blocks and 10 ‘which’ blocks. Study #2 successfully replicated all the main results of study #1 in the ‘confidence’ blocks, we therefore pooled both data sets. In total, we have 81.520 individual decisions and 2565 confidence reports. We have reported all measures, conditions and data exclusions.

### Computational modeling of choice and confidence

#### Bayesian knowledge update

From a bayesian point of view, observers begin each block with a prior distribution *Beta(ps,pf)* for the reward probability of both machines, where *ps* and *pf* encode fictitious prior successes and failures, respectively. A natural reparameterization of this distribution is by using its mean *b=ps/(ps+pf)*, which is a prior measure of the expected payoff, and *w=ps+pf*, which encodes the weight of prior evidence (noted as *Beta(b,w)*). Note that there is a one-to-one mapping between both parametrization, for example, a value of *w*=20 and *b*=0.6 corresponds to a value of *ps*=12 and *pf*=8. Intuitively, low values of the prior mean *b* correspond to a pessimistic perspective, while high values of this parameter represent a more optimistic take. The value of *w* represents how much data it is needed to change the initial prior, higher prior weights require more incoming data in order to change. Due to the conjugacy between the beta prior and the binomial likelihood assumed for the rewards, the posterior distribution after observing *s* successes and *f* failures results in a *Beta(ps+s,pf+f)*.

#### Computation of statistical confidence

Given the posterior distribution *Beta(ps+s,pf+f)* at the moment of the report, the normative statistical confidence that the agent should report after deciding (using any decision process) that machine B has a higher payoff that machine A is:

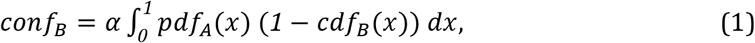

where *pdf_A_(x)* is the value of the Beta density function of machine A evaluated in *x* and *1 — cdf_B_ (x)* is the proportion of the Beta density function of machine B that lies in values higher than *x*. The normalization constant *α* assures that *conf_B_* + *conf_A_ = 1*.

Intuitively, the process of computing confidence in the decision that machine B pays more than A can be seen as taking an infinite number of samples from A, and for each sample calculating the proportion of the distribution of B that lies over it, or equivalently, the probability of taking a sample from B that is bigger than a sample from A (see Fig. 4a). Importantly, the confidence that the agent should report on her decision depends only on the posterior beliefs distribution at the moment of the report and the decision made: it does not depend on how the decision was made, or which was the gameplay strategy that led to the posterior at the moment of the report.

Conceivably, the probability of being correct depends on both the perceived difficulty (equation (2)), and the precision of the distributions at the moment of the report. In practice, however, confidence levels computed with equation (1) are by far dominated by the perceived difficulty. This happens because the confidence report was always implemented at the same time step in the block, so the precision did not vary much between different blocks. Therefore, for building an intuition on the results of Figures 4 and 5 we focused on how the perceived difficulty is different for agents with different priors, but omitted the analyses of how the precision of the final distributions may differ between these agents. Note, however, that the model computes confidence from the posterior belief distribution with the correct precision and perceived difficulty, and these are the model results we show in Figs. 4, 5 and 6. In other words, while the intuitions we built for ‘why’ does confidence biases occur does not account for the precision, the model *does* account for the precision.

#### Decision-making model

Computing the probability of having made a correct decision is straightforward. By contrast, the optimal solution for deciding which machine to play given the experience so far in order to maximize future rewards in finite-horizon bandit problems is more challenging. However, this is a well studied problem that has been solved by dynamic programming-looking at all possible outcomes from the last trial to the current one-, an approach that requires an amount of calculations that grows exponentially with the number of remaining trials [52, 53].

Several heuristics have been developed in order to approximate the optimal solution or mimic human judgements [53, 62]. In this study, the decision of which machine to play is modeled as follows. First, we define the variable *d* (perceived difficulty) as one minus the absolute difference between the means of both machines’ posterior distribution:

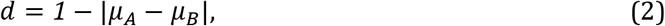

where *μ_A_* and *μ_B_* are computed using the agents’ prior belief (see Eq. 4). A decision value is then sampled from a normal distribution with mean *1-d* (equal to |*μ_A_ – μ_A_* |) and standard deviation σ which we set equal to 0.05 throughout. If the sample is negative, then the machine with the lower estimated payoff is chosen. If it is positive, the arm with the higher estimated payoff is chosen. The probability of sampling a positive value from this distribution, if *μ_A_ > μ_B_* is given by:

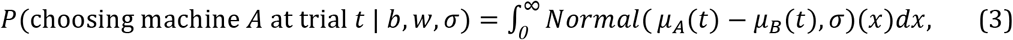

where *Normal*(*μ_A_*(*t*) – *μ_B_* (*t*), <)(*x*) is the Normal density function with mean *μ_A_*(*t*) – *μ_B_* (*t*) and standard deviation *σ* evaluated at point *x*; and *μ_A_* is the mean payoff of machine *A* at trial *t* given the prior bias *b=ps/ps+pf*, the weight *w=ps+pf* and the number of successes (*s_k_*) and failures (*f_A_*) in machine A up to trial *t*:

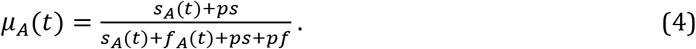

#### Alternative decision-making models

Several arguments support the choice of this heuristic as a model for the decision-making component when compared to other alternatives, like the Win-Stay-Lose-Shift (WSLS) model, the ε-greedy model (which choose the machine with the higher payoff so far with probability 1-ε), and the optimal model, which computes which machine to play by dynamic programming. First, this heuristic captures the proportion of times the option with the lower payoff is chosen as a function of the perceived and unbiased difficulty of the task, which is not the case for most other heuristics (like ε-greedy, which predicts a constant function) or the optimal model. Second, as shown in Fig. 2, it captures the number of times humans persist on playing a machine that is not giving rewards (which is equal to zero for the WSLS model). Third, by accounting solely for *d*, this heuristic ignores the uncertainty about the estimated reward rates (*w*) and assumes a fixed randomness of choice across individuals (σ), so that all the variation in the behavior of different participants can be accounted for only by a different prior mean *b* (Fig. 2). This assumption agrees with the statistical analysis of the data, which shows that the model that varies only *b* was a better model when compared to any other model with one, two or three degrees of freedom (xp>0.99).

### Statistical Analyses

#### Maximum likelihood fitting

We fitted each participant’s prior beliefs with MLE using their decisions in the gameplay. Specifically, given each value of *b* (prior bias), *w* (prior bias weight) and *σ*, we can assign, under our gameplay model, a probability to each decision made by the participant using Eq. 3. Assuming the data is i.i.d., the probability of all the decision from all blocks for a certain participant, given the values of *b, w* and *σ* is:

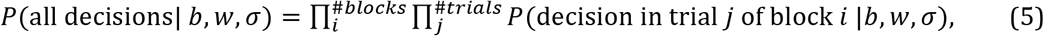

where each term is calculated using Eq. 3. The MLE approach is to choose, for each participant, the set of values {*b, w, σ*} that maximizes the likelihood function (Eq. 5). We evaluated the likelihood function using a discrete grid with 10 uniformly distributed points between 0 and 1 for the parameter *b*, 10 uniformly distributed points between 1 and 150 for *w*, and 8 logarithmically distributed points between *e*^-5^ and *e* for *σ*. When collapsing the data from all participants, the ML values are *w*=20, *σ*=0.05 and *b*=0.5.

#### Model comparison

The exceedance probability (xp) of a model quantifies the probability that it is more frequent than the others (within the tested set of models) in the general population of subjects [63]. We computed exceedance probabilities from the model evidence values of all subjects and all models using the software developed by Stephan et al [63]. The set of competing models are the 7 different models that vary any combination of the parameters *b, w*, and *σ* (the model that varies only *b*, the model that varies *b* and *w*, and so on). The model evidence for a given participant *p(y|M)* quantifies the likelihood of the participant’s data *y* under the model *M*, by integrating over its free parameters θ:

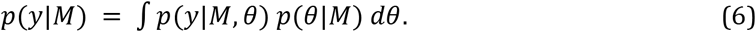

Under the i.i.d data assumption, the probability of making decisions *y_116_* in each block given the randomness of choice σ, the prior value about the machine payoff *σ*, and the weight of this prior *w*, is:

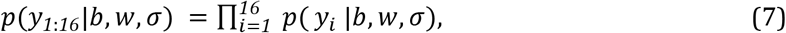

where each term in the product is calculated using Eq. 3. The probability of the decisions from all blocks is simply the product of this last expression over all blocks. Finally, to compute the model evidence for each model *M* we just integrate over its free parameters, and set all the bound parameters to their global MLE value (*w* to 20, *b* to 0.5 and *σ* to 0.05). For example, the model M_k_ with *b* as the only free parameter has the following model evidence:

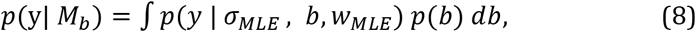

where *p(b)* is uniform and therefore can be omitted. In practice, the integral was approximated by a sum over a discrete grid. Grid points (10 points for *b* and *w*, 8 for σ) were spaced linearly for parameters *b* and *w* and exponentially for σ (which corresponds to a non-informative prior in log-space, a natural choice for variance parameters). As parameter *b* is bounded between 0 and 1, but *w* and σ are unbounded, the final result can depend on the integration limits. Limits were chosen so that the behavior of the model did not change significantly for parameter values exceeding them (*w* between 2 and 150; σ between *e*^-5^ and *e^2^*). Similar exceedance probability values are obtained when the limit was moved plus or minus two points in the chosen scale for each parameter.

#### Two-dimensional presentation of the results

Isoconfidence lines were computed using triangulation-based natural neighbor interpolation (MATLAB *natural* method in the *griddata* interpolation function). Note that not all regions in the difficulty-generosity space shown in Figs. 5b and 6 are allowed, for instance, the block cannot be easy if both machines pay very little.

#### Regression analyses

Were performed using MATLAB *fitlm* function in the Statistical Toolbox. Some of the adjusted values for participants were not reported in the main text. For the function 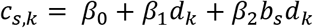, we obtained: 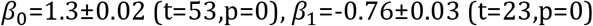.

#### Data availability

The behavioural data and experimental task are available at: https://figshare.com/articles/Behavioral_data/4788823

## Acknowledgements

We would like to thank Falk Lieder, Tania Lombrozo and Tom Griffiths for insightful conversations, Melisa Bentivegna for help with early data collection, and the organizers of the Subjective Confidence Workshop at Les Treilles where some preliminary results were first presented. This work was partly funded by grants CONICET PIP 11220130100384CO and UBACYT 20020130200202BA. FM is funded by the French center for atomic energy (CEA).

